# IGF2 Peptide-Based LYTACs for Targeted Degradation of Extracellular and Transmembrane Proteins

**DOI:** 10.1101/2023.10.30.563730

**Authors:** Michał Mikitiuk, Jan Barczyński, Przemysław Bielski, Marcelino Arciniega, Urszula Błaszkiewicz, Aleksandra Hec, Andrea D. Lipińska, Michał Rychłowski, Tad A. Holak, Tomasz Sitar

## Abstract

Lysosome Targeting Chimeras (LYTACs) have recently been developed to facilitate lysosomal degradation of specific extracellular and transmembrane molecular targets. However, the LYTAC particles described to date are based on glycopeptide conjugates, which are difficult to prepare and produce on a large scale. Here we report the development of pure protein LYTACs based on the non-glycosylated IGF2 peptides, which can be readily produced in virtually any facility capable of monoclonal antibody production. These chimeras utilize the IGF2R/CI-M6PR pathway for lysosomal shuttling and, in our illustrative example, target programmed death ligand 1 (PD-L1), eliciting physiological effects analogous to immune checkpoint blockade. Results from in vitro assays significantly exceed the effects of anti-PD-L1 antibodies alone.

## Introduction

Lysosome-targeting chimeras (LYTACs) represent a new technology that recruits extracellular and/or membrane proteins for their degradation via the endosome/lysosome pathway^1-6^. LYTACs, also called MoDEs for molecular degraders of extracellular proteins^3^, are bifunctional conjugates that simultaneously bind the extracellular domain of a target protein of interest (POI) and a cell-surface lysosome-targeting receptor (LTR) to form a ternary complex, leading to protein internalization via clathrin-mediated endocytosis^7^ and subsequent degradation in a lysosome. Currently known LYTACs at the POI-targeting side contain antibodies, small molecules, peptides, aptamers^1-6,8-17^ These are linked to the LRK-binding glycopeptides, peptides, aptamers, dendritic DNA, and cytokines that bind to the receptors facilitating endocytosis and lysosomal degradation: the cation-independent mannose-6-phosphate receptor (CI-M6PR),^1, 11-14^, the liver-specific asialoglycoprotein receptor^3,4,9,15^, integrin^8^, the transmembrane E3 ligase ring finger 43 (RNF43)^10^, surface scavenger receptors,^16^ the cytokine decoy recycling receptor CXCR7^17^.

The first lysosomal internalization receptor used for LYTACs was the CI-M6PR, also known as the insulin-like growth factor 2 receptor (IGF2R). The CI-MPR/IGFR2 is a type I transmembrane glycoprotein of approximately 300 kDa^18^. The extracellular region of the CI-MPR has 15 homologous domains (124–192 amino acids each). CI-M6PR mainly binds mannose 6-phosphate (M6P)-bearing proteins. The M6P-binding sites are located in domains 3, 5, 9, and 15^18,19^. The acidic pH of the lysosome triggers the release of the glycosylated cargo for degradation by the lysosomal enzymes and acid hydrolases. The receptor is then shuttled back to the membrane to repeat the cycle^20,21^.

CI-MPR/IGFR2, in contrast to the cation dependent MPR (CD-MPR), also binds a number of non-glycosylated ligands, such as, for example, the non-glycosylated insulin-like growth factor 2 (IGF2)^22-27^. The binding of IGF2 is mediated by the IGF2-binding domain 11 of CI-MPR/IGFR2^18,19^. It is thought that when the CI-MPR/IGFR2 is present on the cell surface, domain 11 binds to any free IGF2 in the extracellular matrix^19^. The receptor is then rapidly internalized together with IGF2 via a YSKV motif present in its cytoplasmic tail^28^.

The LTR-binding ligands for CI-MPR are the mannose-6-phosphonate (M6Pn)-derived glycopolypeptides (PolyM6Pn) whose synthesis is complicated. It starts with the conversion of mannose pentaacetate to M6Pn-NCA in 13 steps and ends with copolymerization of M6Pn-NCA to give poly(M6Pn) polypeptides^1^. Another drawback is that the conjugation of this synthetic poly(M6Pn) ligand to serine or lysine residues on antibodies results in a complex inhomogeneity of conjugate structures^1,2,29^.

Herein, we report the development of a series of structurally well-defined IGF2-peptides that can be incorporated to and expressed with antibodies targeting proteins of interest and successfully internalize and degrade these proteins via the IGF2R/CI-M6PR pathway (**Fig. 1**). This will greatly facilitate the development of IGF2R/CI-M6P-based LYTACs for therapeutic applications.

**Fig. 1.**
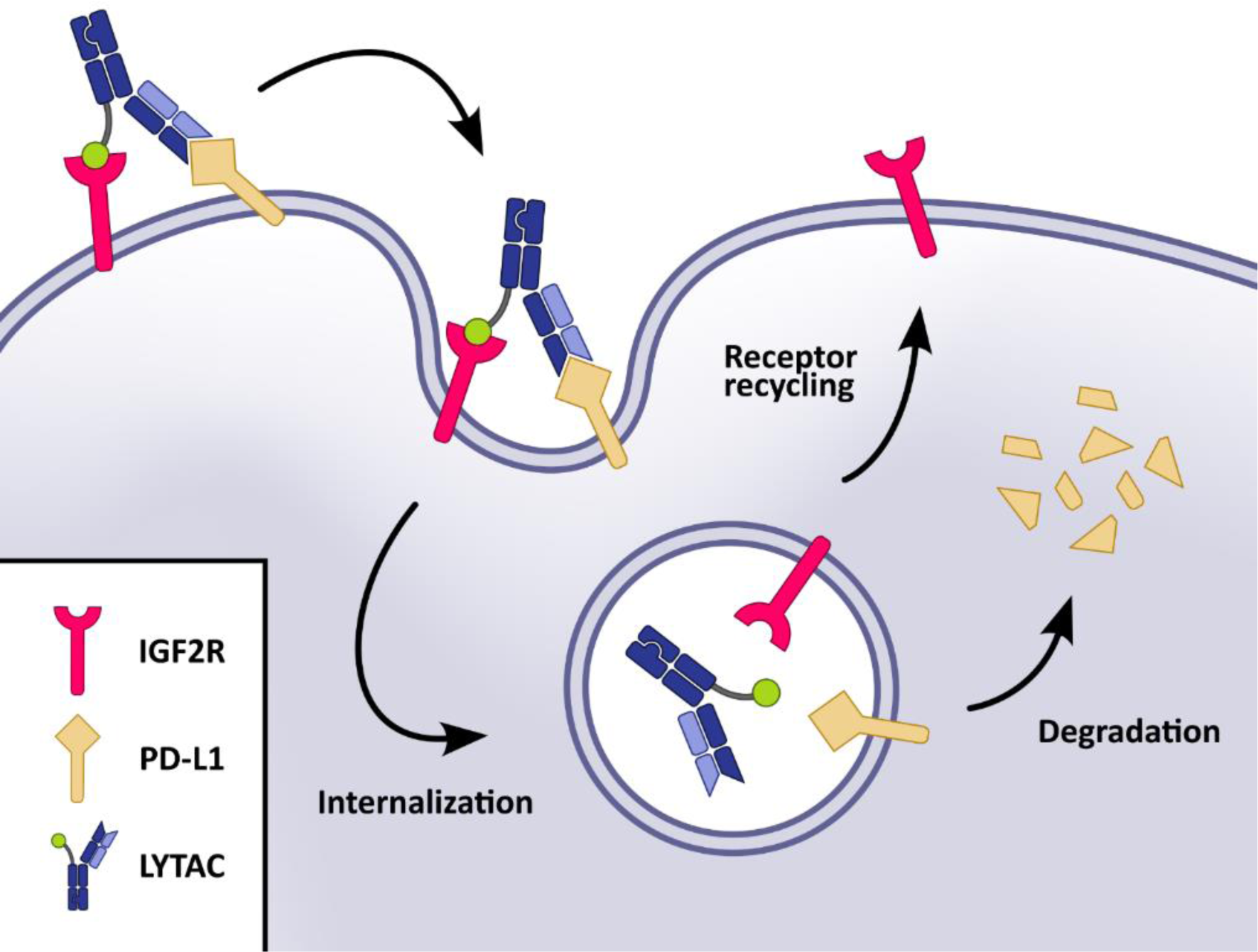
Schematic representation of the targeted transmembrane PD-L1 degradation through the IGF2R/CI-M6PR pathway. The procedure uses the IGF2-peptide based and protein-only LYTAC compound.

## Results

### The designed IGF2-based polypeptides bind to IGF2R but not to IGF1R

We designed and produced IGF2-based peptides, which were then used to design the PD-L1-degrading bispecific LYTAC compounds (Supplementary Table 1). These IGF2 polypeptides bind to domain 11 of the IGF2R with different affinities, but all show higher affinities to this domain than wild-type IGF2 (Fig. 2a). In addition, all generated polypeptides showed virtually no binding to IGF1R. This is in contrast to the wild-type IGF2, which also showed binding to IGF1R (Fig. 2b).

**Fig. 2.**
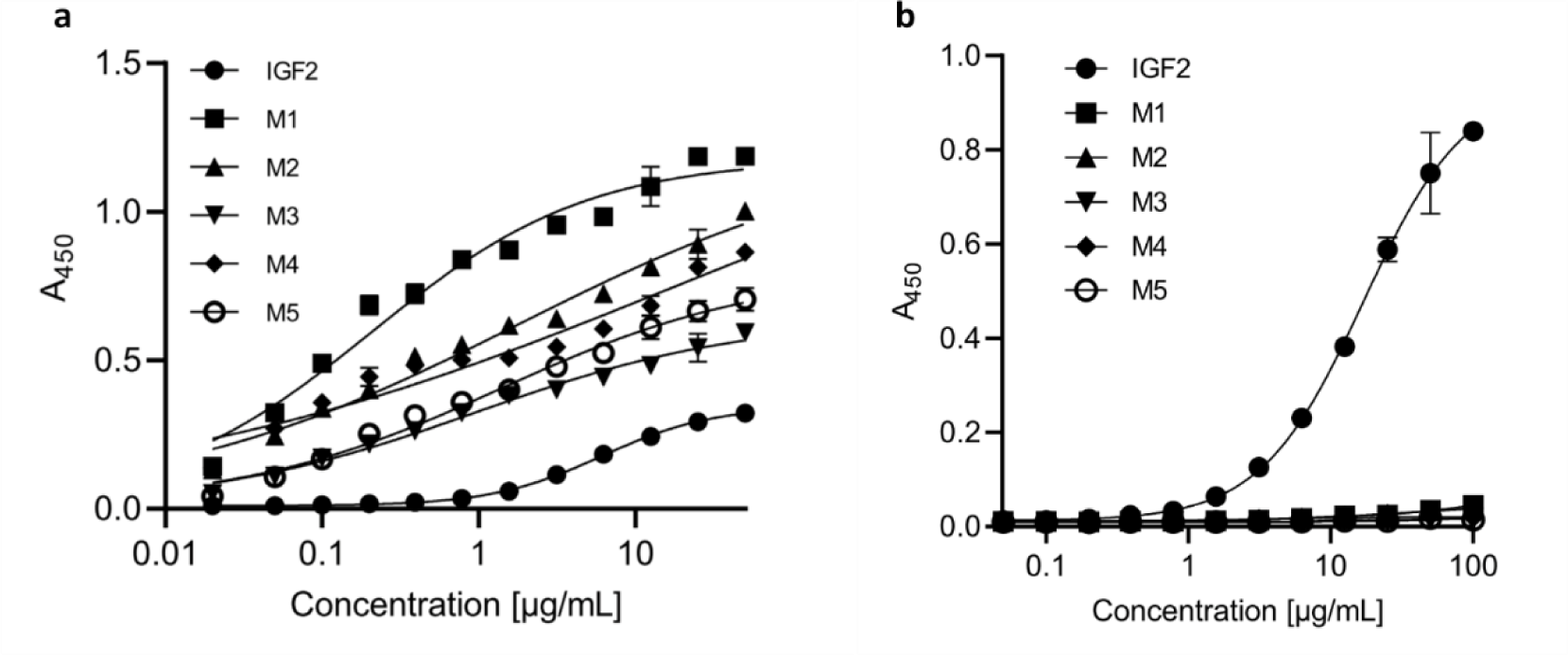
All generated IGF2-like peptides bind IGF2R-D11 with different affinities, whereas they do not bind to IGF1R. **a** Assessment of binding to domain 11 of IGF2R by ELISA. **b** Assessment of binding to IGF1R by ELISA.

### Design, production, and characterization of bispecific anti-PD-L1-anti-IGF2R compounds

Based on the results of the binding assays, the polypeptide labeled M1 in **Fig. 2** was selected for the construction of a bispecific molecule along with previously modeled and produced recombinant human anti-PD-L1 antibody (Supplementary Table 2). The antibody, designated C5, bound PD-L1 with a slightly higher affinity than the reference anti-hPD-L1 antibody durvalumab (**Fig. 3a**). However, C5 was not able to disrupt the PD-1/PD-L1 binding on its own (**Fig. 3b**).

**Fig. 3.**
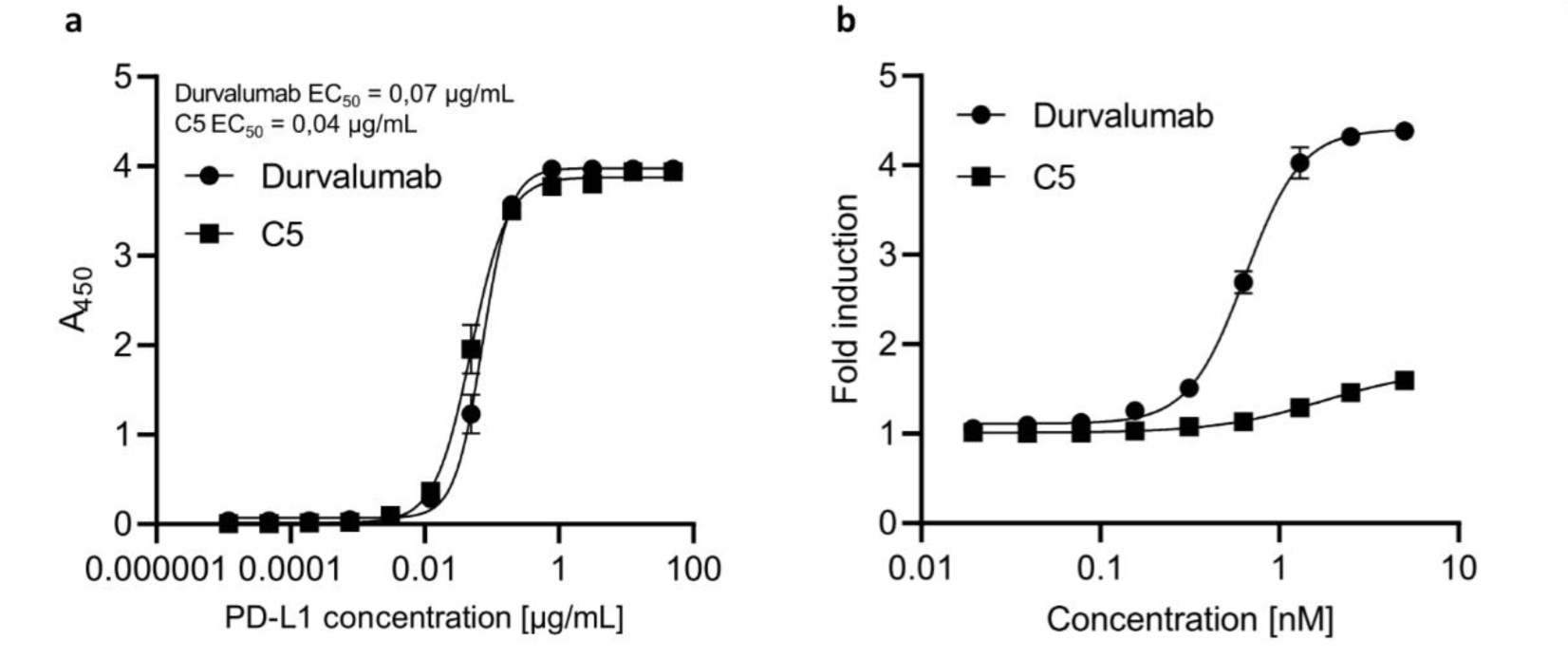
Comparison of durvalumab and C5 antibodies. **a** Binding of antibodies to PD-L1 by ELISA. **b** Ability to disrupt PD-1/PD-L1 binding in the PD-1/PD-L1 blockade bioassay.

As the final selection of the bispecific protein format requires in vitro and in vivo functional characterization^30^, we designed the LYTAC bispecific molecules in two formats for early screening: as asymmetric knob-into-hole (KIH) C5 IgG4 with one of the Fabs replaced by M1 (designated as C5M1A) and C5 scFv with M1 attached to its C-terminus by a (GGGGS)_4_ linker (designated as C5M1B) (Fig. 4a). The constructs were prepared and purified as described in the Methods. Their ability to bind to appropriate targets was confirmed by ELISA (Fig. 4b and Fig. 4c). As expected, both bispecifics bound to both IGF2R and PD-L1, whereas IGF1R was not bound by either. C5M1B showed a higher affinity for PD-L1 overall and for IGF2R at lower concentrations. However, a higher signal was observed for C5M1A at higher concentrations.

**Fig. 4.**
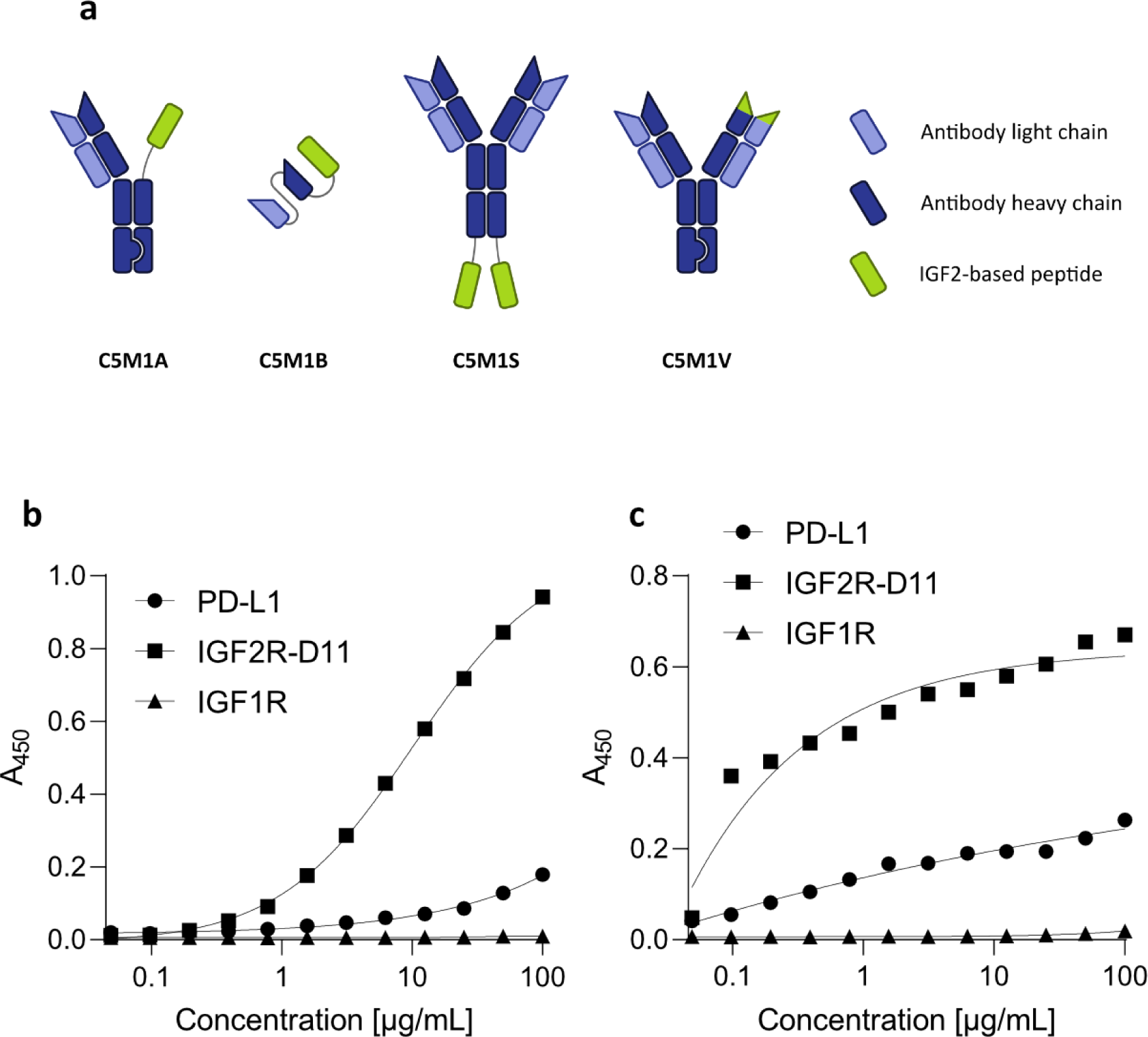
Produced bispecific LYTAC compounds. **a** Schematic representation of selected formats (C5M1A, C5M1B) and formats planned for production (C5M1S, C5M1V). **b** Results of C5M1A binding to PD-L1, IGF2R-D11 and IGF1R determined by ELISA. **c** Results of C5M1B binding.

### The bispecifics induce internalization of soluble PD-L1

Having confirmed that both compounds bind specific targets, we tested their ability to internalize PD-L1. Initial tests using a microplate reader showed a statistically significant result for C5M1A compared to the untreated cells. The controls tested (C5 antibody and M1 IgG1 Fc peptide) did not yield statistically significant results when compared to untreated cells (Supplementary Fig. 1).

To further confirm that the soluble proteins were properly internalized, live fluorescence microscopy experiments were performed. Both compounds C5M1A and C5M1B were able to induce internalization of the soluble PD-L1-mCherry (Fig. 5). Furthermore, the observed internalization occurred in a time-dependent manner (Supplementary Fig. 2, Supplementary Fig. 3). C5M1B showed a lower rate of internalization than C5M1A.

**Fig. 5.**
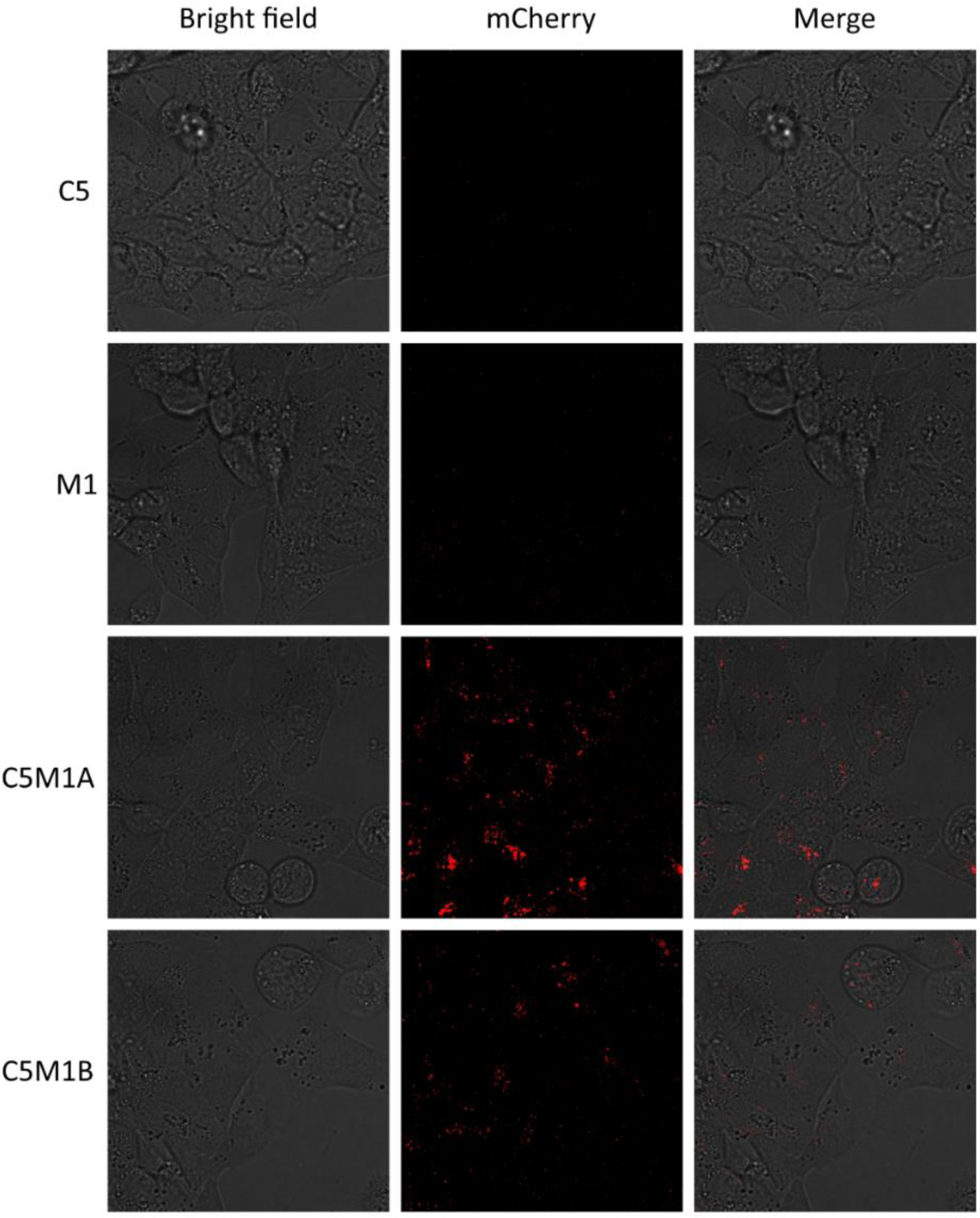
Fluorescence microscopy images of RL95-2 cells treated with the control (C5, M1-IgG1Fc chimera) and LYTAC (C5M1A, C5M1B) compounds at 100 nM with 100 nM od PD-L1-mCherry fusion protein for 22 h.

### Bispecific compounds induce internalization of transmembrane PD-L1

To confirm whether transmembrane proteins are internalized, flow cytometry experiments were performed. After 22 h of incubation with the bispecific compounds, the Panc 10.05 cell line showed a significant reduction of PD-L1 on the cell surface (Fig. 6a). Similar to all previously described LYTAC compounds, the molecular target was degraded in a concentration-dependent manner (Fig. 6b). In contrast to the ELISA results, C5M1A induced a higher degradation (54.9%) than C5M1B (38.9%) at the same concentration. Correspondingly, the RL95-2 cell line showed a significant level of surface PD-L1 degradation after C5M1A treatment (Supplementary Fig. 4), further confirming that the method can target different tissues.

**Fig. 6.**
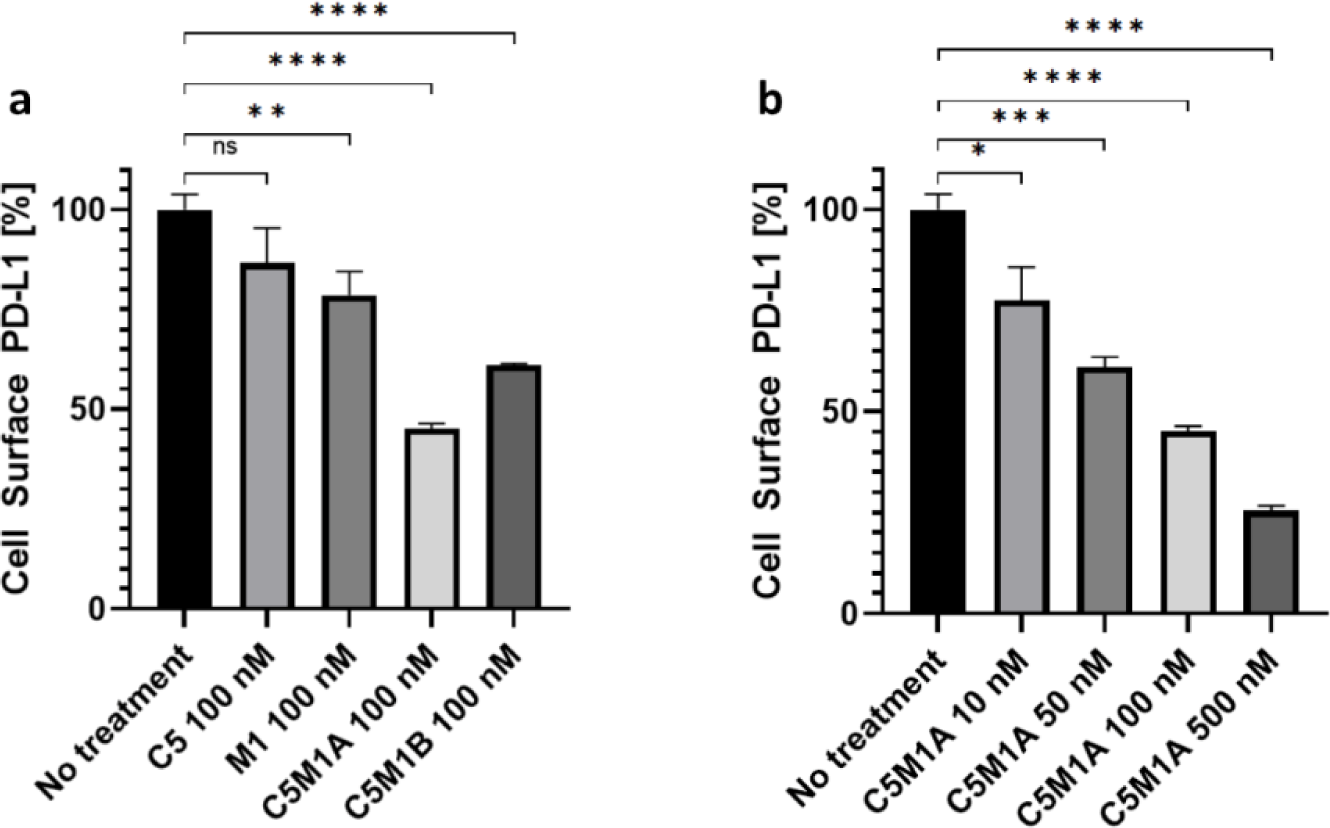
Determination of cell surface PD-L1 levels by live cell flow cytometry. **a** Panc 10.05 cells treated with bispecific LYTAC and control compounds. **b** Panc 10.05 cells treated with rising concentrations of C5M1A. Data on all charts represent mean from 3 independent replicates after background signal subtraction as mean ± SD. Untreated control was considered baseline level (100%). The unpaired t-test was used to compare the means of each group against untreated control. P value threshold of less than 0,05 was considered statistically significant.

### Bispecific compounds induce tumor cell cytotoxicity when incubated with PBMC

After establishing the internalization potential of the produced LYTAC compounds, their efficacy was evaluated in a peripheral blood mononuclear cell (PBMC) cytotoxicity assay. Incubation of PBMC with tumor cells and immune checkpoint degraders results in significantly higher levels of growth inhibition/lysis than controls (C5 antibody and M1-IgG1Fc peptide alone) (Fig. 7). The intensity of growth inhibition depends on a number of factors, including the tumor cell line used and the abundance of PD-L1 on the cell surface, the format of the LYTAC molecule, its concentration, and the activity of the cells isolated from the donor. While the activity of the isolated cells varied, both compounds were tested in a meaningful number of different donors. In order to minimize the variability resulting from testing in multiple donors, C5M1A was tested at various concentrations in PBMC from a single donor (Supplementary Fig. 5a), which also resulted in a significant level of tumor cell lysis compared to the control compound. To further confirm applicability of the method in varying tissues, both bispecific compounds were tested on BT20 cell line, yielding 38% and 26,6% of growth inhibition for C5M1A and C5M1B, respectively (Supplementary Fig. 5b).

**Fig. 7.**
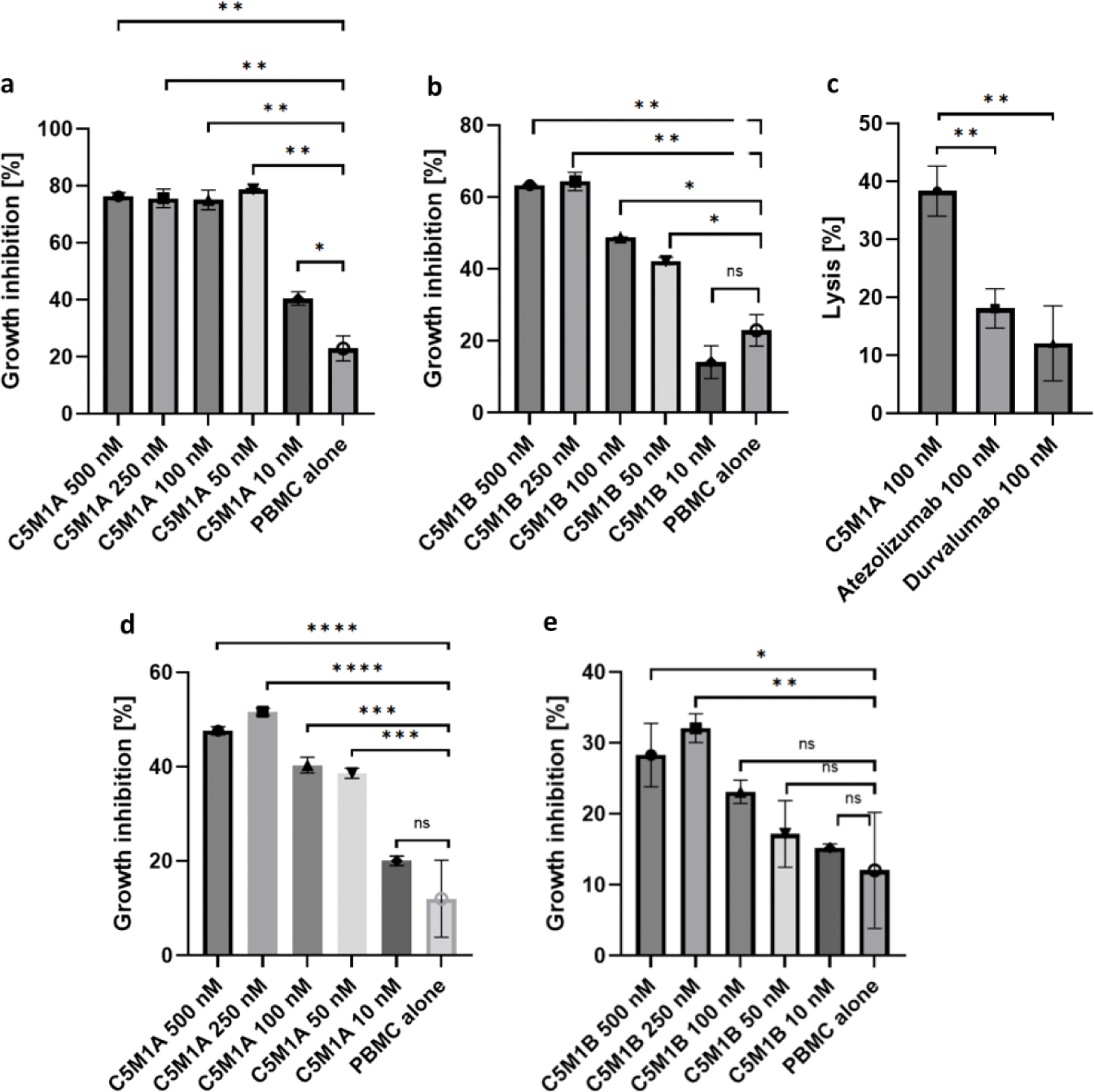
Results of PBMC cytotoxicity assays of the two LYTAC compounds. a RL95-2 cells treated with C5M1A; **b** RL95-2 cells treated with C5M1B; **c** Efficiency of lysis induced by C5M1A, atezolizumab and durvalumab on RL95-2 cells; **d** PANC-1 cells treated with C5M1A; **e** PANC-1 cells treated with C5M1B; each concentration was tested on PBMC from different donors. Data in all graphs represent the mean of 3 independent replicates as mean ± SD. The unpaired t-test was used to compare the means of each group with the untreated control. P-value threshold of less than 0.05 was considered statistically significant.

Results regarding compounds activity obtained from both, live fluorescence microscopy and flow cytometry are further confirmed: the asymmetric format induces higher growth inhibition than scFv. Efficacy is also dependent on cell line type: RL95-2 was more sensitive to growth inhibition than PANC-1.

Finally, to rule out potential cytotoxicity of the synthesized compounds alone, a cytotoxicity test for C5M1A was carried out. The results showed minimal cytotoxicity from the compound solution alone (Supplementary Fig. 6).

## Discussion

Extracellular and membrane-associated proteins comprise approximately 40% of the human proteome^33^ and are key players in cancer, age-related diseases, and autoimmune disorders^34^. Bispecific molecules that direct the internalization and degradation of these proteins have shown great promise as a therapeutic approach.^1-16^ While such molecules have been reported for some time,^35-37^ it is only recently that specific endosomal targeting molecules such as LYTAC, KineTAC, etc. have been described in the literature.^1-16^

LYTACs are sugar-conjugated antibodies that bind the target protein and a lysosome-targeting receptor, i.e., cation-independent mannose-6-phosphate receptor (CI-M6PR)^1,5^ or asialoglycoprotein receptor (ASGPR),^2-5^ thereby inducing internalization, lysosomal trafficking, and degradation of membrane proteins. The LYTAC compounds described to date are based on conjugates, which would be challenging for large-scale production and therefore not feasible as potential therapeutic agents. In this study, we developed novel protein-only bispecific LYTACs capable of inducing soluble and transmembrane PD-L1 internalization in a time- and concentration-dependent manner. Since no chemical modifications are required to produce such compounds, their large-scale production can be easily adopted by virtually any monoclonal antibody manufacturing facility. Another desirable feature that further streamlines the manufacturing process is that our IgG-based bispecific molecule does not cause the light chain mismatching problem that is common in IgG bispecific formats because it contains Fab on only one arm. Finally, genetically encoded compounds have the potential to be delivered as mRNAs or as part of adoptive cell therapies.

Two recent methods for membrane protein degradation are bispecific antibody-based PROTACs (AbTACs)^10^ and cytokine receptor-targeting chimeras (KineTACs).^17^ These bispecifics have one part that binds to the target protein and the other part that binds to a transmembrane E3 ligase (AbTACs: RNF43) or cytokine receptor (KineTACs: CXCR7). Our work differs from LYTAC and the latter two approaches in the use of a short peptide rather than a glycopolymer (LYTAC) or a bispecific antibody with the whole cytokine as in the case of KineTAC.

Our protocol is somewhat similar to that of the aptamer-based LYTACs. However, aptamer-based LYTACs are challenged by their rather unfavorable pharmacokinetics, such as rapid clearance from the blood. It is also necessary to ensure their resistance to nucleases present in serum^31,32^. They also require chemical modifications, which makes the manufacturing process complicated. Finally, only one aptamer-based therapeutic has been approved for human use to date, so such an approach may face additional challenges during the regulatory approval process. As a result, the development of aptamer-based drugs is associated with higher risk.

From the results of the C5 antibody analysis, it is clear that to construct functional LYTACs, the specificity-determining moiety does not have to block the interaction between the ligand and the corresponding receptor. This eliminates the step of selecting antibody candidates based on their ability to disrupt a specific binding, further streamlining the process of generating new, potentially therapeutic entities. Furthermore, the use of a non-antagonist antibody as the POI targeting arm suggests that the observed cytotoxicity is due to target degradation rather than binding disruption.

Comparison of the two formats shows that the knob-into-hole IgG construct induces a higher fold of degradation compared to the smaller scFv-peptide fusion, despite the use of long, flexible linkers to combine all elements of the latter construct. This may indicate that the binding geometry provided by the KIH IgG construct is more favorable for facilitating degradation. These results highlight the importance of proper format selection, as the asymmetric construct induced a significantly higher rate of internalization than the scFv. However, all variables, such as serum half-life, must be considered when selecting a format for the bispecific.

A desirable feature of our designed LYTAC compounds (although based on IGF2) is that they do not bind to the IGF1R. This allows us to avoid potential unwanted side effects of IGF1R overstimulation. Protein-only LYTACs are able to induce cell surface protein degradation at nanomolar concentrations. The effect obtained in our experiments appears to exceed the efficacy of currently available monoclonal antibodies targeting PD-L1 at the same concentration in vitro. The development of bispecific degraders may be associated with a hook effect, where only one arm of the compound remains bound to its target, reducing degradation efficiency and requiring higher doses. Concentrations tested up to 500 nM do not appear to reduce efficiency - in fact, degradation efficiency increases at higher concentrations.

The results obtained are consistent with previously described conclusions that LYTACs utilizing CI-M6PR/IGF2R can be applied to a wide range of tissues^1^, however, the efficiency of degradation appears to depend on the amount of the target protein on the cell surface of a specific tissue cell.

Our data demonstrate the potency of the IGF2-based degraders and identify them as excellent candidates for further preclinical evaluation.

## Methods

### Chemicals and reagents

All bispecific compounds, anti-PD-L1 antibodies, including reference antibodies: IgG4 durvalumab and atezolizumab, IGF2-based polypeptides, PD-L1-mCherry fusion protein (R1-002-5), IGF2, domain 11 of IGF2R and IGF1R were produced by Recepton, except for antibodies used for detection in ELISA and flow cytometry. Genetic constructs were synthesized by Eurofins Genomics. Plasmid DNA midiprep (cat. K210004) was purchased from Thermo Fisher Scientific. All restriction enzymes and T4 ligase (cat. M0202L) were purchased from New England Biolabs. Culture media for CHO (cat. 94120) and HEK (cat. 9413) were purchased from Fujifilm Irvine Scientific. L-glutamine (cat. HN08.3) was purchased from Carl Roth. RPMI-1640 (cat. 30-2001), Dulbecco’s Modified Eagle’s Medium (DMEM) (cat. 30-2002), DMEM: F-12 (cat. 30-2006) and Eagle’s Minimum Essential Medium (EMEM) (cat. 30-2003) were purchased from ATCC. Trypsin (cat. 25200-072) was purchased from Thermo Fisher Scientific. Fetal bovine serum (cat. P30-19375), hygromycin B (cat. P06-08100) and DPBS (cat. P04-361000) were purchased from PAN Biotech. G418 (geneticin, cat. G073-39US) was purchased from TOKU-E. Polyethylenimine linear (25 kDa) (cat. 23966-100) was purchased from Polysciences. Mouse anti-human PD-L1 antibody (cat. 14-5983-82) and goat anti-mouse IgG Alexa Fluor 488 conjugated antibody (cat. A11001) were purchased from Thermo Fisher Scientific. Goat anti-mouse Ig PE-conjugated antibody (cat. 550589) was purchased from BD Biosciences. Insulin (cat. I9278) was purchased from Sigma. Bovine serum albumin (cat. BP9702-100) was purchased from Fisher Scientific. 10x Phosphate Buffered Saline was made with 80 g NaCl (cat. 27810.295, VWR); 2,0 g of KCl (cat. 0395, VWR); 14,4 g of Na_2_HPO_4_ (cat. 117992300, Chempur); 2,4 g of KH_2_PO_4_ (cat. 26925.295, WVR) and pH adjusted to 7,4. It was then diluted 1:10 to achieve 1x working solution. PBST was made by adding 0,5% Tween 20 (cat. M147, VWR) to the 1x PBS. PD-1/PD-L1 blockade bioassay (cat. J1250) was purchased from Promega.

### Design of IGF2-based polypeptides with abrogated IGF1R binding

An experimental structure of human IGF2 was used as a template for the design (6UM2.pdb). Specifically, the template consisted of segments Glu6-Ser29 and Gly41-Pro63. The 12-residue gap between Ser29 and Gly41 was filled with shorter linker sequences (2 to 3 residue length) using the Rosetta Suite^38^. First, the experimental template was subjected to a relax protocol^39^ with optimization of hydrogen bonds and sidechain amide group. Rotamers from the input structure were used in packing, in addition to extra sampling for chi1 and ch2 rotamers. Backbone coordinates were tether of the initial template coordinates. The maximum number of minimization cycles was set to 200. A single structure was requested as an output. Based on the relaxed structure two runs of the remodel protocol^40^ were performed considering linkers of length 2 and 3, respectively. The number of requested structures was set to 500 for each run. After a structural inspection of top results (according to the Rosetta score), 5 models were selected for experimental evaluation.

### Stable cell lines generation

For production of bispecific compounds, IGF2, IGF2-based polypeptides and IGF1R stable cell lines were generated. Genetic constructs were cloned into expression plasmids, linearized and cells were transfected by electroporation, followed by selection with increasing concentrations of G418 (geneticin) and/or hygromycin B. Productivity of each generated cell line was assessed by Ultra-Performance Liquid Chromatogrpahy (UPLC), cells with satisfactory productivity were propagated and 1 L batch of each protein was produced. In case of IGF1R, obtained cell pool has undergone clonal selection to further increase productivity.

### Transient protein production in HEK293

For production of domain 11 of IGF2R and PD-L1-mCherry, transient production in HEK293 has been utilized. Obtained genetic constructs were cloned into expression plasmids. Cells were seeded at 10^6^ cells/mL and transfected by addition of DNA-PEI (1:3) solution, using 1 ug DNA per 1 mL of medium. Cells were incubated at 37 °C, 180 rpm, 8% CO_2_ in a S41i incubator (Eppendorf) for 5 days.

### Purification of proteins

All cultures on the day of the harvest were centrifuged and the supernatant was filtered through 0,2 µm polyethersulfone filter (Advanced Microdevices). Purification of the proteins was performed by affinity chromatography and size exclusion chromatography (SEC). Filtrates containing PD-L1-mCherry and C5M1B proteins were passed through an equilibrated (20 mM NaHPO_4_ + 300 mM NaCl + 5 mM imidazole pH 7.4) Ni Sepharose Excell (Cytiva) column, then the column was washed with 20 mM NaHPO_4_ + 300 mM NaCl + 10 mM imidazole pH 7.0. Protein was eluted with 20 mM NaHPO_4_ + 300 mM NaCl + 500 mM imidazole pH 7.0 using step gradient (protein eluted at 50% B). PD-L1-mCherry was loaded on the equilibrated Superdex 200 Increase column (Cytiva) and the main peak was pooled.

All other proteins were either antibodies or they contained the IgG1 Fc tag, added for purification purposes. Filtrates were passed through an equilibrated (20 mM NaHPO_4_ + 150 mM NaCl pH 7.0) MabSelect Sure (Cytiva) column, then the column was washed with the same buffer. Protein was eluted with 100 mM citric acid pH 3.3. The eluate pH was adjusted to 6.8. Then, all purified proteins except C5M1A, were loaded on equilibrated (PBS) Superdex 200 Increase column (Cytiva) and the main peak was pooled.

### ELISA

ELISA plates (Nunc MaxiSorp, Thermo Fisher) were coated with 50 µL solution of proteins at the appropriate concentration and left overnight (at 4°C). The next day, plates were equilibrated at RT, washed (4 x 300 µL PBST) and blocked for 1h at RT with 1% BSA (Fisher Scientific) in PBS solution. Ligands were diluted in PBS to desired concentrations, added to appropriate wells (50 µL) and left for 1h incubation at RT. Next, plates were washed (4 x 300 µL PBST) and primary antibodies were added in 1 : 10000 dilution (anti-IgG Fc cat. 31789, Thermo Fisher; anti-His-Tag cat. 4603-08, SouthernBiotech) followed by 1 h incubation at RT. After another wash, HRP-conjugated streptavidin was added (cat. 21124, Thermo Fisher) in 1 : 10000 dilution (50 µL) and incubated for 1 h at RT. Finally, plates were washed (6 x 300 µL PBST) and 100 µL of pre-warmed 3,3’,5,5’ tetramethylbenzidine (TMB) solution (Merck-Sigma) was added to each well. The assay was developed for 6 min, followed by adding stop solution (0,2 M H_2_SO_4_). Absorbance at 450 nm (655 nm background substraction) was read on Tecan Spark microplate reader, data were analyzed in GraphPad Prism software (v9.5.1) using log(agonist) vs. response – variable slope (four parameters) model.

### Fluorescence internalization test

Human endometrium carcinoma RL95-2 cells (ATCC CRL-1671) were seeded in 96-well plates at 50 000 cells/well. After 24h the medium was exchanged and individual compounds (100 nM) along with PD-L1-mCherry fusion (100 nM) in PBS were added. Cells were incubated at 37°C for another 22 h, after which the medium was discarded, cells were washed twice with PBS, and fluorescence of each well was measured using Tecan Spark microplate reader with excitation at 590 nm and emission at 620 nm.

### Live fluorescence microscopy

RL95-2 cells were seeded in 10-well glass bottom plates to reach 50% confluency on the day of the observation. The medium was exchanged and compounds (100 nM) along with PD-L1-mCherry fusion protein (100 nM) in PBS were added. Specimens were imaged for 22 h using a confocal laser scanning microscope (Leica SP8X equipped with an incubation chamber for the live analysis) with a 63× oil immersion lens (Leica, Germany). Excitation 585 nm, emission 602 nm – 651 nm (red-mCherry). LAS X software was used for data analysis.

### Flow cytometry

Human pancreatic adenocarcinoma Panc 10.05 cells (ATCC CRL-2547) were cultured in 6-well plate for 24 h at 0,6·10^6^ cells/well. The medium was exchanged and compounds at appropriate concentrations in PBS were added. After 22 h of incubation cells were trypsynized and 0,4·10^6^ cells were collected in 1,5 mL tubes. Cells were centrifuged at 1300 g for 3 min and incubated in cold 0,5% BSA-PBS solution containing mouse anti-PD-L1 antibody (1:50) and incubated on ice for 60 min. Next cells were washed twice with cold 0,5% BSA-PBS and subsequently incubated with Alexa Fluor 488- or PE-modified secondary antibody (diluted 1:1000 or 1:350, respectively). For background signal control, cells were incubated only with secondary antibodies. Cells were washed twice with cold 0,5% BSA-PBS and resuspended in 0,5% BSA-PBS. 10 000 cells were analyzed using BD FACSCalibur cytometer. Cell Quest Pro software was used for data analysis.

### PD-1/PD-L1 blockade bioassay

The PD-1/PD-L1 immune checkpoint bioassay (PD-1/PD-L1 Bioassay, Promega) was performed according to the manufacturer’s manual. PD-L1+ aAPC/CHO-K1 cells were plated in 96-well, white, flat bottom assay plates at 40x104 cells in 100 µL of medium (Ham’s F12, 10% FBS) and incubated overnight at 37°C, 5% CO2. The next day medium was removed from the assay plate and serially diluted antibodies added at 40 µL per well in the assay buffer (RPMI1640 + 1% FBS + 1% DMSO). Next, PD-1 Effector Jurkat cells (included in the assay kit) were resuspended in assay buffer (RPMI 1640 + 1% FBS) at a concentration of 1,25 x 106 cells/mL and added to the assay plate at 40 µL per well (total of 50 x 104 cells). The cells were co-cultured for 6 h (37°C, 5% CO2) and then removed from incubator and equilibrated at room temperature for 5 min. Bio-GloTM Reagent (Promega) was prepared according to the manufacturer’s manual and added to each well at 80 µL per well. Assay plates were incubated in room temperature for 15 min, luminescence was measured on the Tecan Spark microplate reader. Data were analysed in GraphPad Prism software (v.9.5.1) using log(inhibitor) vs. response – Variable slope (four parameters) model.

### Human tumor cells killing assay

Peripheral blood mononuclear cells (PBMC) were isolated through Ficoll gradient centrifugation (Ficoll Paque Plus, Cytiva) from human blood samples from healthy individuals (obtained from the Regional Centre for Blood Donation and Treatment in Gdańsk, Poland). After isolation, cells were cryopreserved in 90% FBS (FBS Good, PANBiotech), 10% DMSO (Sigma Aldrich). A day prior the experiment human tumor cells were plated on 96-well plate (3x10^4^ cells/well) in an appropriate growth medium. Simultaneously, PBMC effectors were thawed and allowed to rest overnight in RPMI 1640 medium (ATCC), 10% FBS (FBS Good, PANBiotech). Various concentrations of compounds were tested with maintained 10 : 1 effector : target (E:T) cells ratio. Assays were incubated for 120 h in 37°C with 5% CO_2_. After this time 20 ul of MTT (5 mg/mL, PanReac AppliChem) was added to each well and left for 2 h incubation, followed by the addition of 100 µL of MTT crystals dissolvent (10% SDS, 0,01 N HCl). Plates were left in the incubator (37°C, 5% CO_2_) overnight. The next day absorbance was read at 570 nm with background subtraction at 690 nm (Tecan Spark). Cell lysis percentage was calculated as a ratio of compounds-treated samples (human tumor cells + PBMC + tested compounds) to PBMC-treated samples (human tumor cells + PBMC). Growth inhibition was calculated as a ratio of treated samples (human tumor cells + PBMC +/- compounds) to non-treated samples (human tumor cells). Statistical analysis was performed in GraphPad Prism software (v.9.5.1).

### Cytotoxicity test

A day prior the experiment human tumor cells were plated on 96-well plate (3x10^4^ cells/well) in an appropriate growth medium. The next day, dilutions of tested compound were prepared and added to tumor cells. Assays were performed for 120 h in the incubator (37°C, 5% CO_2_). After incubation 20 µL of MTT (5 mg/mL, PanReac AppliChem) was added to each well and left for 2 h incubation, followed by addition of 100 µL MTT crystals dissolvent (10% SDS, 0.01 N HCl). Plates were left in the incubator (37°C, 5% CO_2_) overnight. The next day absorbance was read at 570 nm with background subtraction at 690 nm (Tecan Spark). Data are presented as a percent of living cells relative to no treatment control.

## Acknowledgements

This research was supported by Grant POIR.01.01.01-00-0129/18 from the National Centre for Research and Development, Poland. M.M, J.B., P.B. and A.H. were supported by Ph.D. studentships awarded by the Ministry of Science and Higher Education, Poland (DWD/4/54/2020 and DWD/3/11/2019).

## Author Contributions

M.M. and J.B. produced proteins and carried out most of the experiments and analyses. P.B. purified the proteins. M.A. designed the IGF2-based peptides. A.D.L. carried out flow cytometry analysis. M.R. carried out live fluorescence microscopy experiments. U.B. and A.H. provided intellectual input. T.S. provided supervision. M.M., J.B., P.B., M.A., T.A.H. and T.S. wrote the manuscript with discussion and feedback from all co-authors.

