## Supplementary information for "IGF2 Peptide-Based LYTACs for Targeted Degradation of Extracellular and Transmembrane Proteins"

### List of Supplementary Tables

Supplementary Table 1 Aminoacid sequences of designed IGF2-based peptides.

Supplementary Table 2 Aminoacid sequence of durvalumab-based anti-hPD-L1 IgG4 antibody C5. The antibody does not display ability to disrupt PD-1/PD-L1 interaction on its own.

### List of Supplementary Figures

Supplementary Fig. 1 Determination of PD-L1-mCherry uptake by RL95-2 cells by fluorescence measurement.

Supplementary Fig. 2 Fluorescence microscopy images of RL95-2 cells treated with the C5M1A.

Supplementary Fig. 3 Fluorescence microscopy images of RL95-2 cells treated with the C5M1B.

Supplementary Fig. 4 Determination of cell surface PD-L1 levels by live cell flow cytometry of RL95-2 cells treated with C5M1A.

Supplementary Fig. 5 Results of PBMC cytotoxicity tests.

Supplementary Fig. 6 Cytotoxicity test of C5M1A.

**Supplementary Table 1 Aminoacid sequences of designed IGF2-based peptides.**

| | |
|---|---|
| M1 | ETLCGGELVDTLQFVCGDRGFYNDDGIVEECCFRSCDLALLETYCATP |
| M2 | ETLCGGELVDTLQFVCGDRGFYKDDGIVEECCFRSCDLALLETYCATP |
| M3 | ETLCGGELVDTLQFVCGDRGFYFSGGIVEECCFRSCDLALLETYCATP |
| M4 | ETLCGGELVDTLQFVCGDRGFDMRGGIVEECCFRSCDLALLETYCATP |
| M5 | ETLCGGELVDTLQFVCGDRGFYFSGIVEECCFRSCDLALLETYCATP |

**Supplementary Table 2 Aminoacid sequence of durvalumab-based anti-hPD-L1 IgG4 antibody C5.** The antibody does not display ability to disrupt PD-1/PD-L1 interaction on its own.

| | |
|---|---|
| Light chain | IVLTQSPGTLSLSPGERATLSCKASEDIGTWLAWYQQKPGQAPRLLIYDASSRA<br>TGIPDRFSGSGSGTDFTLTISRLEPEDFAVYYCQQYGSLPWTFGQGTKVEIKRT<br>VAAPSVFIFPPSDEQLKSGTASVVCLLNNFYPREAKVQWKVDNALQSGNSQESV<br>TEQDSKDSTYSLSSTLTLSKADYEKHKVYACEVTHQGLSSPVTKSFNRGEC |
| Heavy chain | EVQLVESGGGLVQPGGSLRLSCAASGFTFSRYWMSWVRQAPGKGLEWVALISRD<br>GSETYYVDSVKGRFTISRDNAKNSLYLQMNSLRAEDTAVYYCAREGGWFGELAF<br>DYWGQGTLVTVSSASTKGPSVFPLAPSSKSTSGGTAALGCLVKDYFPEPVTVSW<br>NSGALTSGVHTFPAVLQSSGLYSLSSVVTVPSSSLGTQTYICNVNHKPSNTKVD<br>KRVEPKSCDKTHTPPCPPCPAPEFLGGPSVFLFPPKPKDTLMISRTPEVTCVVV<br>DVSQEDPEVQFNWYVDGVEVHNAKTKPREEQFNSTYRVVSVLTVLHQDWLNGKE<br>YKCKVSNKGLPSSIEKTISKAKGQPREPQVYTLPPSQEEMTKNQVSLTCLVKGF<br>YPSDIAVEWESNGQPENNYKTTPPVLDSDGSFFLYSRLTVDKSRWQEGNVFSCS<br>VMHEALHNHYTQKSLSLSLGK |

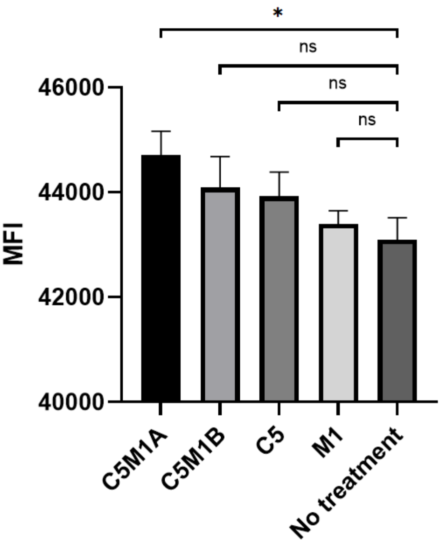


**Supplementary Fig. 1 Determination of PD-L1-mCherry uptake by RL95-2 cells by fluorescence measurement.** All compounds added at 100 nM, treated for 22 h. Data on all charts represent mean from 3 independent replicates as mean ± SD. The unpaired t-test was used to compare the means of each group against untreated control. P value threshold of less than 0,05 was considered statistically significant.

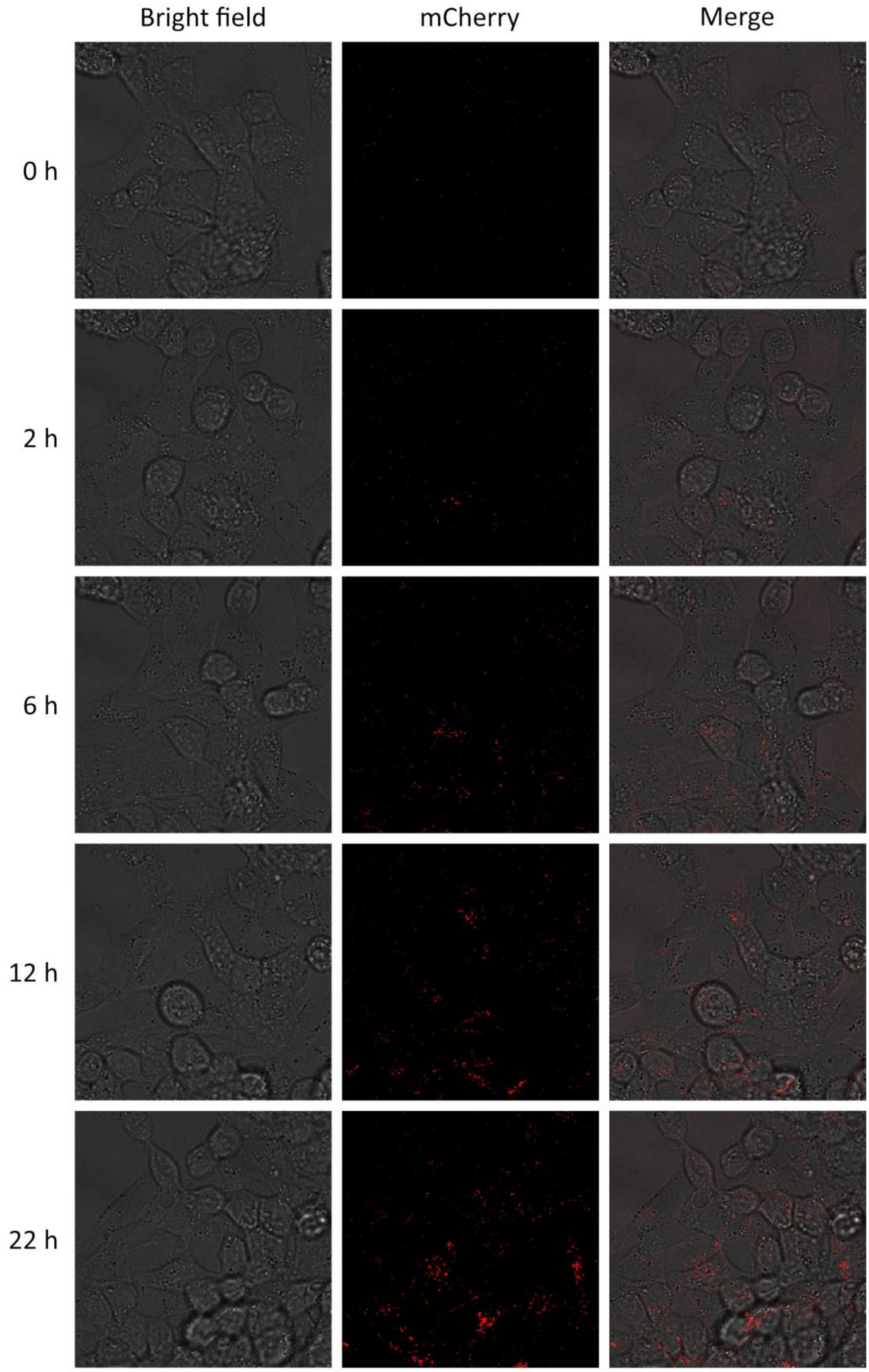


**Supplementary Fig. 2 Fluorescence microscopy images of RL95-2 cells treated with the C5M1A** at 100 nM with 100 nM of PD-L1-mCherry fusion protein over 22 hours.

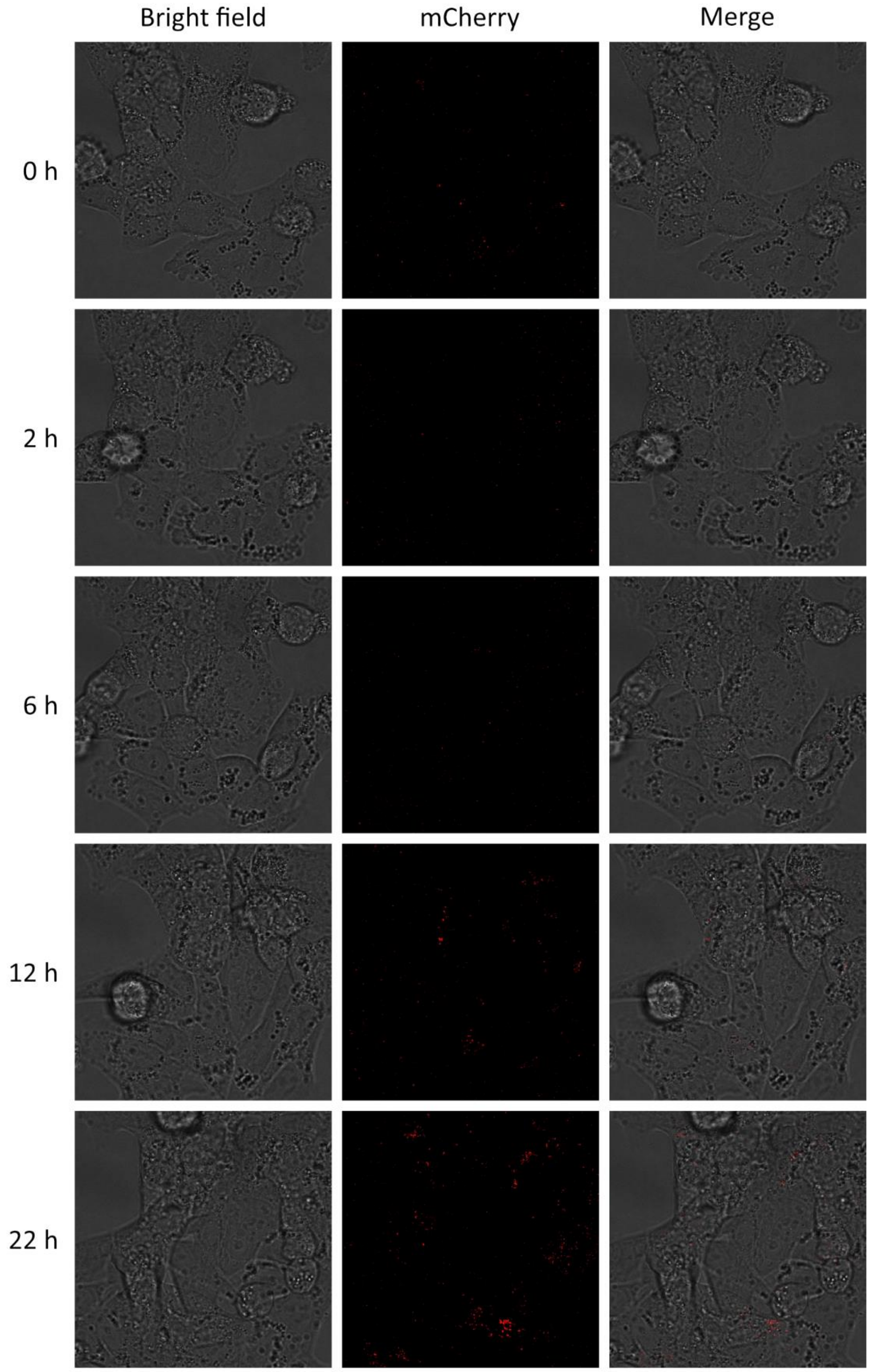


**Supplementary Fig. 3 Fluorescence microscopy images of RL95-2 cells treated with the C5M1B** at 100 nM with 100 nM of PD-L1-mCherry fusion protein over 22 hours.

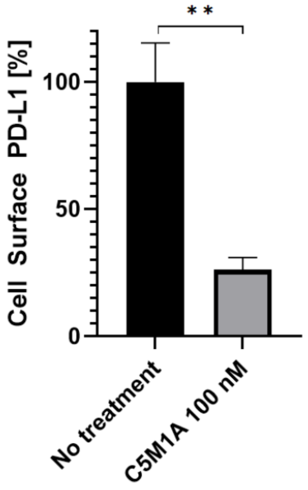


**Supplementary Fig. 4 Determination of cell surface PD-L1 levels by live cell flow cytometry of RL95-2 cells treated with C5M1A.** Data represent mean from 3 independent replicates after background signal substraction as mean ± SD. Untreated control was considered baseline level (100%). The unpaired t-test was used to compare mean of experimental group against untreated control. P value threshold of less than 0,05 was considered statistically significant.

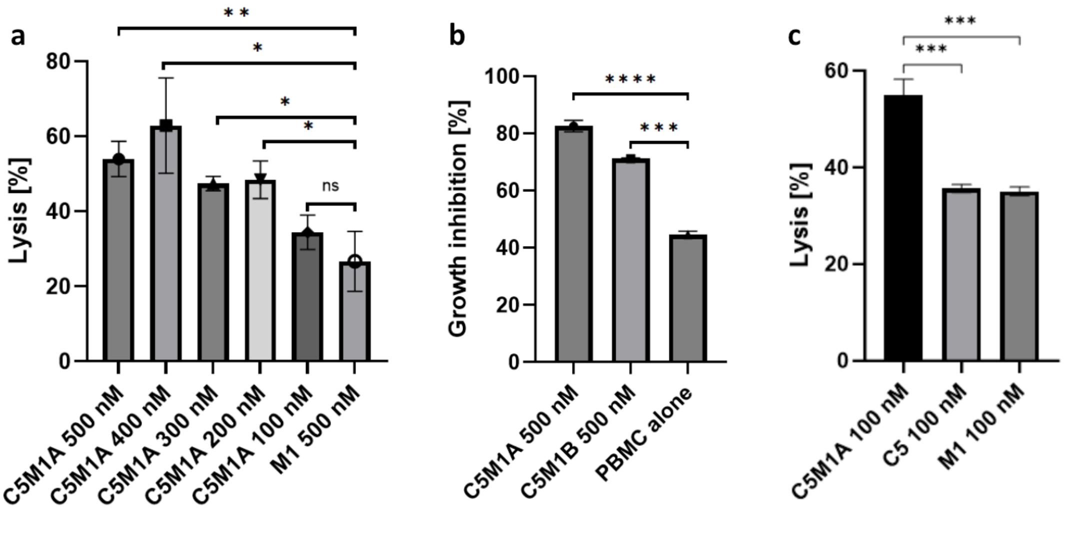


**Supplementary Fig. 5 Results of PBMC cytotoxicity tests. a** RL95-2 cells treated with C5M1A tested with PBMC from a single donor, compared to 500 nM M1-IgG1Fc chimera; **b** BT20 cells treated with C5M1A and C5M1B. **c** RL95-2 cells treated with C5M1A and control compounds: M1-IgG1Fc chimera and full C5 antibody. Data on all charts represent mean from 3 independent replicates as mean ± SD. The unpaired t-test was used to compare the means of each group against untreated control. P value threshold of less than 0,05 was considered statistically significant.

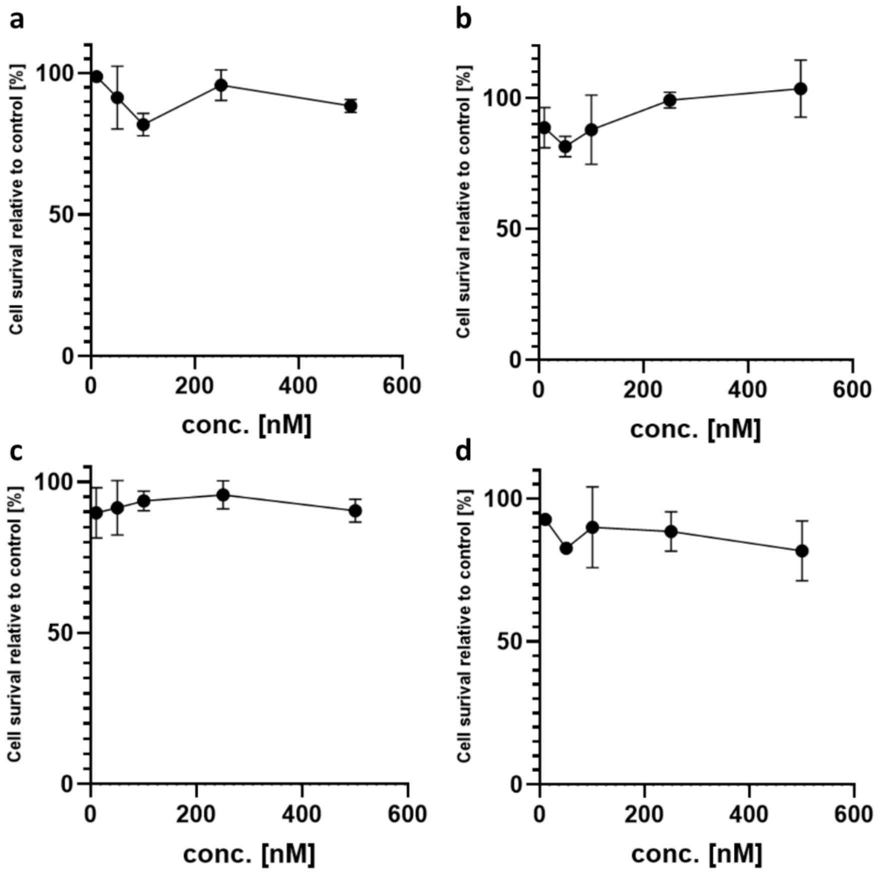


**Supplementary Fig. 6 Cytotoxicity test of C5M1A.** All used cell lines were analysed: a RL95-2 b Panc 10.05 c PANC-1 d BT20
